# Self-assembled DNA nanostructures promote cell migration & differentiation of human umbilical vein endothelial cells

**DOI:** 10.1101/2022.01.28.478149

**Authors:** Anjali Rajwar Gada, Payal Vaswani, Dhiraj Bhatia

## Abstract

DNA nanostructures have been explored for capabilities to influence cellular behavior and its functions. Recent times have seen the development of new emergent functionalities of DNA nanodevices as class of biomaterials with immense capacity to interface with biological systems and having vast potential in disease diagnosis and therapeutics. Being chemically robust and biocompatible in nature, DNA nanostructures have been surface modified and structurally fine-tuned to find emerging applications in the field of stem cell therapy and tissue regeneration. DNA nanostructures can be utilized for therapeutic angiogenesis that involves induction of blood vessel formation and can be used to treat ischemic diseases like stroke or heart failure. This work addresses the effect of DNA nanostructures’ structural topology in their capacity to stimulate endothelial cells angiogenesis. We tested a panel of four geometries of DNA nanostructure and checked their potential on the differentiation of human umbilical vein endothelial cells (HUVECs). While different DNA nanostructure geometries showed successful angiogenesis induction and cell migration in HUVECs, tetrahedral DNA cages showed the maximum uptake and angiogenesis potential indicating that not only the composition of materials, but also the 3D arrangement of ligands might also play role in stimulating the angiogenesis process.

## Introduction

DNA nanotechnology has recently come into focus as a powerful technique that can be utilized to design well-defined structures in 1D, 2D, and 3D with varied shapes and sizes at the nanoscale level^1^. These nanostructures have tunable architectures and offer inherent biocompatibility, ability to encapsulate cargoes, and chemical flexibility, allowing easy conjugation of DNA nanostructures with different peptides, proteins, oligonucleotides, and small molecules etc^2–5^. These fascinating properties of DNA make it an exciting biomaterial that has shown application in diagnosis, drug delivery, and recently in stem cells differentiation and tissue engineering^6,7^. DNA nanostructures have been utilized to trigger proliferation and differentiation of stem cells^8^. Stem cell differentiation is a process that involves the changes in gene expression in parent cells leading to subsequent changes in cell properties and morphology thereby altering their sensitivity to different stimuli leading to the acquisition of modified functions^9^. Different types of stem cells like mesenchymal, embryonic stem cells etc. can be utilized for tissue regeneration^10^. Previous research has shown the potential of DNA nanostructures, specifically tetrahedral DNA (TD), in the proliferation and differentiation of stem cells. When exposed to different populations of stem cells, TD can induce differentiation. DNA TD promoted self-renewal by activating the *Wnt/*β*-catenin* pathway and differentiation via inhibiting the Notch pathway in neuroectodermal stem cells (NE-4C)^8^. In periodontal ligament stem cells (PDLSCs) DNA Tetrahedron promoted proliferation and induced osteogenic differentiation via *Wnt/*β*-catenin* signaling pathway^11^. DNA nanotube functionalized with peptide can differentiate neural stem cells into neurons^12^.

Angiogenesis is a process of forming new blood vessels from the existing ones via the proliferation, migration, and differentiation of endothelial cells^13^. Angiogenesis plays a key role in cancer metastasis that help in maintaining oxygen and nutrient supply thereby supporting tumor growth^14^. DNA tetrahedron have shown the potential to induce proliferation, migration and differentiation of endothelial cells via notch signaling pathway^14^. These works have encouraged the potential of DNA nanostructure in field of tissue engineering and regenerative medicine. However, the effects of topology of DNA nanostructures on angiogenesis have not been explored in details. Here, we have investigated the effects of different geometries of DNA nanocages (DNs) in the angiogenesis of endothelial cells and explored their potential for tissue engineering. We first checked the uptake of DNs in the endothelial cells and found that tetrahedral DNA showed maximum internalization among all the geometries of DNs tested. We next reviewed the migration and angiogenic potential of DNs via tubule formation and scratch assays. The results showed that enhanced DNs promote cell migration and tubule formation, whereas geometry-specific biases were not observed in the differentiation process. We have also tested different matrices, checked their effect on angiogenesis, and found that DNs are compatible with all the matrices for cell growth and can retain their potential to trigger tubule formation and cell migration. Taken together, this work can establish ground rules for future investigations involving DNA nanocages for biological and biomedical applications, explicitly applying their surface topologies in the areas of bioimaging, drug delivery, immune activation, and tissue engineering.

## Results

### Synthesis and characterization of DNA nanostructures (DNS)

DNA nanocages of different topologies viz. tetrahedron (TD)^15^, icosahedron (ID)^16^, cube^17^ and Bucky ball (BB)^18^ were self-assembled using programmed thermal annealing method. All the DNA nanostructures were assembled using the previously used protocol^19^. The DNs were fluorophore labelled using cyanine-3 modified strand, which was incorporated during the reaction. The assembled products were characterized using electrophoretic mobility shift assay that showed the formation of higher order structure due to the lag in their migration as a function of cage like structure formation.

### Cellular uptake of DNS in HUVECs

DNs needs to bind and internalized by the cells in order to perform the targeted function… Cellular uptake of Cy3 labelled DNs in HUVECs was studied using laser scanning confocal microscopy (LSCM). HUVECS are endothelial cells derived from the human umbilical vein endothelium and are an extensively used as in-vitro model to study angiogenesis. These are primary cells that have shown application in vascular modeling, oncology, pharmacology and tissue engineering^20^. To study the internalization of DNs, HUVECs were incubated with DNs for 20 min in serum free media along with transferrin-A488, which is used as an endocytic marker for clathrin mediated endocytosis. Green channel indicates Tf vesicles and small punctate like structures in the red channel marks the internalization of DNA **(Figure 2a)**. The cellular uptake of DNs was quantified by measuring the intensity of Cy3 DNs in red channel. We observed that number of punctates were more in BB and TD followed by Cube and ID. The more intensity in BB is due to more number of fluorophores in BB than TD. However, the number of fluorophore is more in cube, but the intensity was less than TD. BB’s raw intensity was more than TD, but when the intensity was normalized to the number of fluorophores per cage, it was less than TD **(Figure 2b)**. The uptake of Tf was almost constant with all the DNs **(Figure 2c)**.

**Figure 1.**
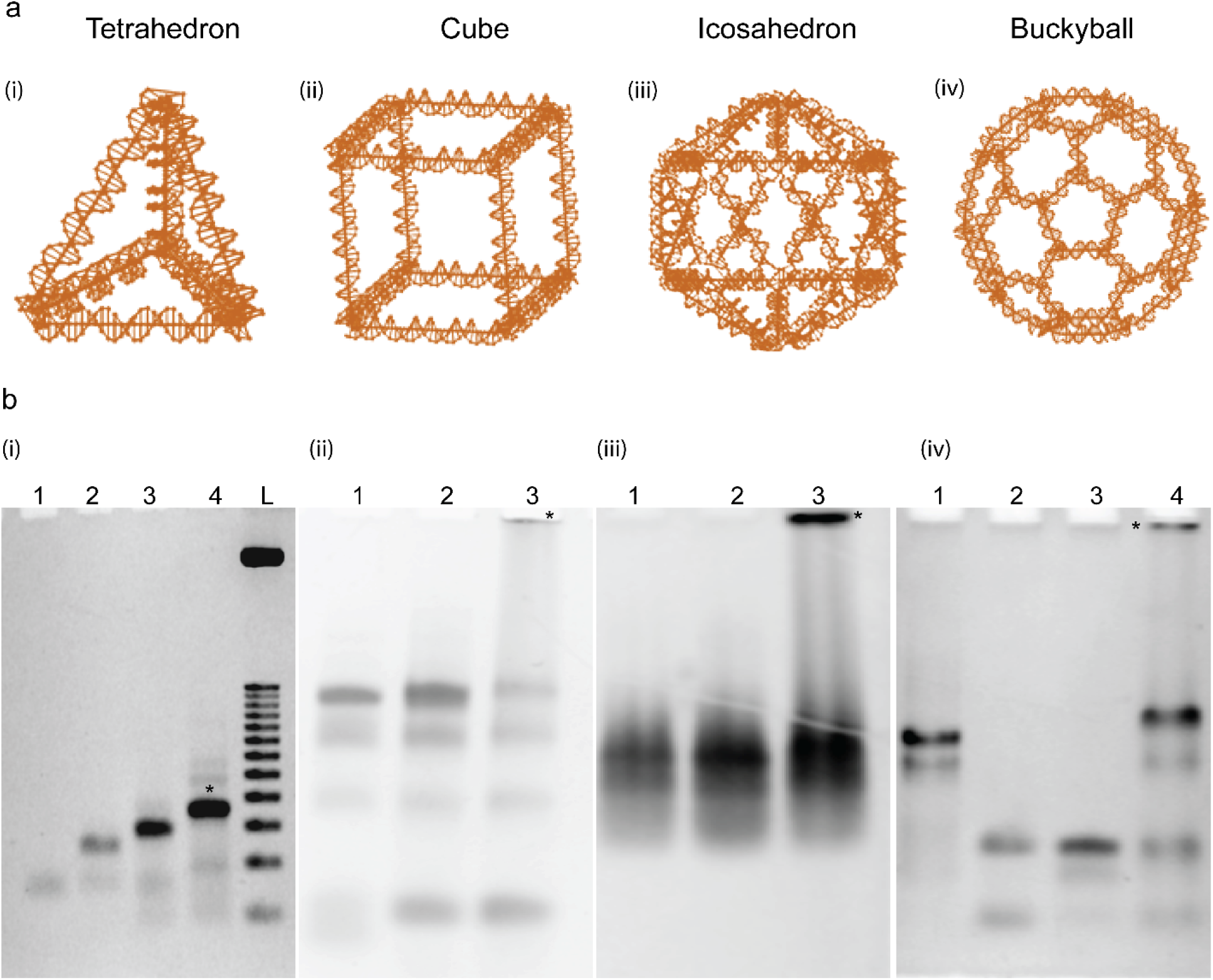
Synthesis and characterization of DNS. **(a)** Nanoengineer design of DNS with different topologies. Tetrahedron (TD), Cube, Icosahedron (ID) and Bucky ball (BB). **(b)** 5% non-denaturing polyacrylamide gel electrophoresis (PAGE). **(i)** Formation of TD from four strands. Lane 1: T1, Lane 2: T1+T2, Lane 3: T1+T2+T3, Lane 4: T1+T2+T3+T4 (TD), L: 100 bp ladder **(ii)** Tile-based assembly of DNA Cube. Lane 1: Tile A, Lane 2: Tile B, and Lane 3: Cube (Tile A+TileB). **(iii)** Modular assembly of ID. Lane 1: VL_5_ (lower half module), Lane 2: VU_5_ (upper half module) Lane 3: Full ID (VU_5_ + VL_5_). **(iv)** Formation of BB from L, M, and S strands. Lane 1: L+M strand, Lane 2: M+S, Lane 3: L+S, and Lane 4: BB (L+M+S). (*) indicates complete structure.

**Figure 2.**
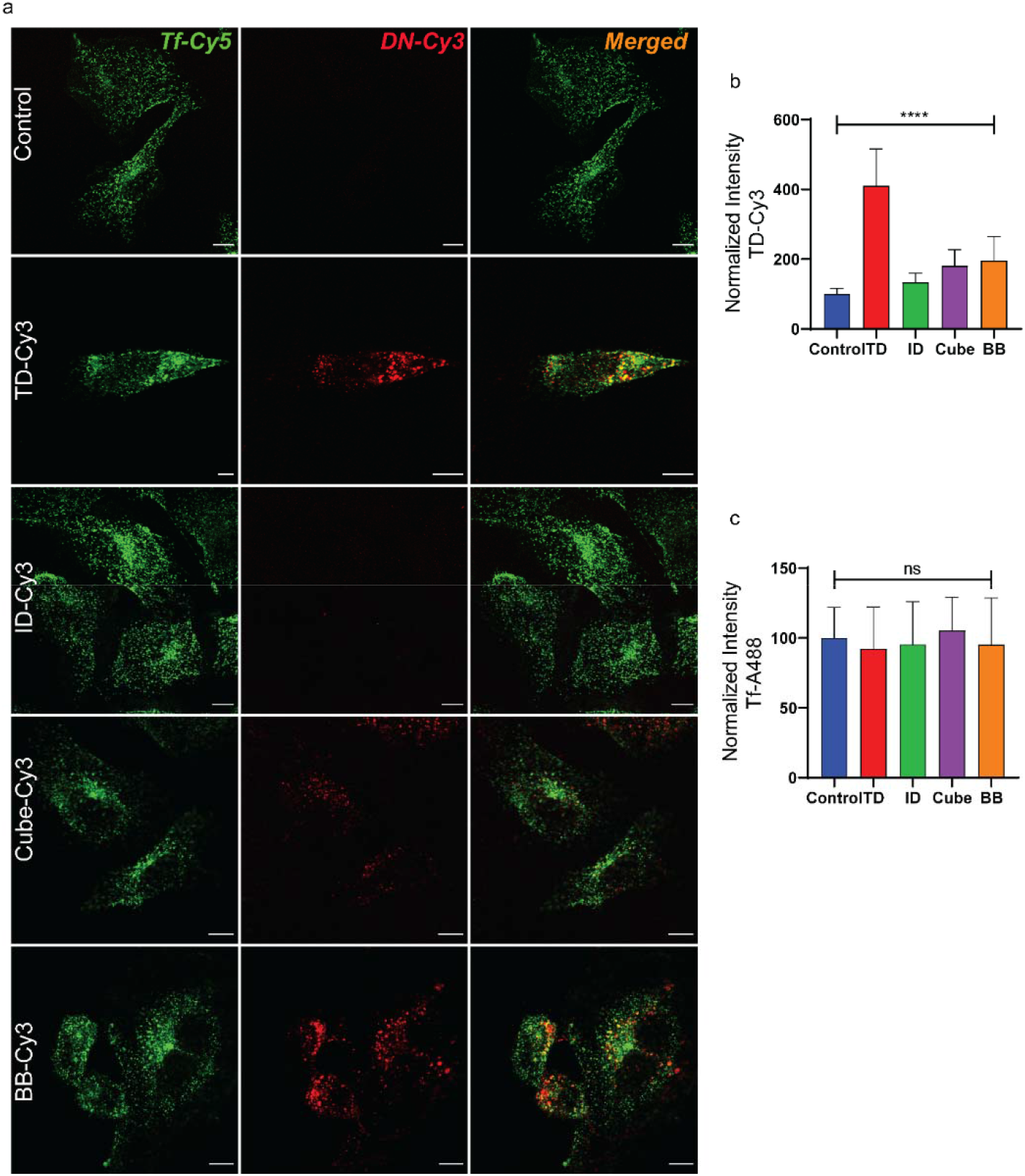
Cellular uptake of DNS in HUVECs. **(a)** LSCM images of endothelial cells (HUVECs) treated with 150 nM of DNs. Green channel indicates the uptake of Tf-Cy5 (5µg/ml) which was kept constant with all the DNs. The red channel indicates the uptake of Cy3 labelled DNs. The last row indicates the merged images of the green and red channel. Scale Bar: 10 µm. **(b)** Quantification of DNs uptake from images in panel (a) Error bars denotes standard errors. The normalized intensity was calculated from 40 cells (ordinary one-way ANOVA, p value < 0.0001****). **(c)** Quantification of Tf-Cy5 uptake from images in panel (a) Error bars denotes standard errors. The normalized intensity was calculated from 40 cells (ordinary one-way ANOVA, ns: non-significant).

### DNA nanostructures stimulate angiogenesis in HUVECs

In order to explore the effects of DNs in angiogenesis, we performed tubule formation assay in human umbilical vein endothelial cells (HUVECs). HUVECs are an extensively studied model for in vitro angiogenesis because they can form tubules like structures similar to capillary in response to different growth factors or suitable stimuli^21^. We first did the concentration-dependent study to check the effect of DNs concentration on tubule formation. We used DNA TD for concentration dependent studies since DNA TD has shown preferential uptake compared to other nanostructures in the previous result. Cells were treated with different concentrations of TD (100 nM – 400 nM) after seeding them on the matrigel-coated plate in EBM media devoid of growth factor and serum for 6 hours. The result showed that DNA TD stimulated the formation of tubular structures. In control where no treatment of DNs was given, the tubulation was less and small capillary-like structures can be seen **(figure 3a)**. There was a substantial increase in the number of junctions and tubule length compared to control at higher concentrations. At 400 nM, the polygonal network of tubules was observed **(figure 3b)**. We have used 400nM for further studies. We next checked the effect of time in the tubule formation. We incubated cells with DNA TD (400 nM) and monitored the tubule formation at different time points. The cells were mostly aggregated with round morphology at time (t = 0 h), and with the time, cells were attaching to form tiny tubules and at t = 3 h, bigger tubules can be observed. The mature tubule-like polygonal network can be seen t = 6 h **(figure 3c)**. The number and size of tubules increased with time, and the entire tubule network was visible at 6 h **(figure 3d)**.

**Figure 3.**
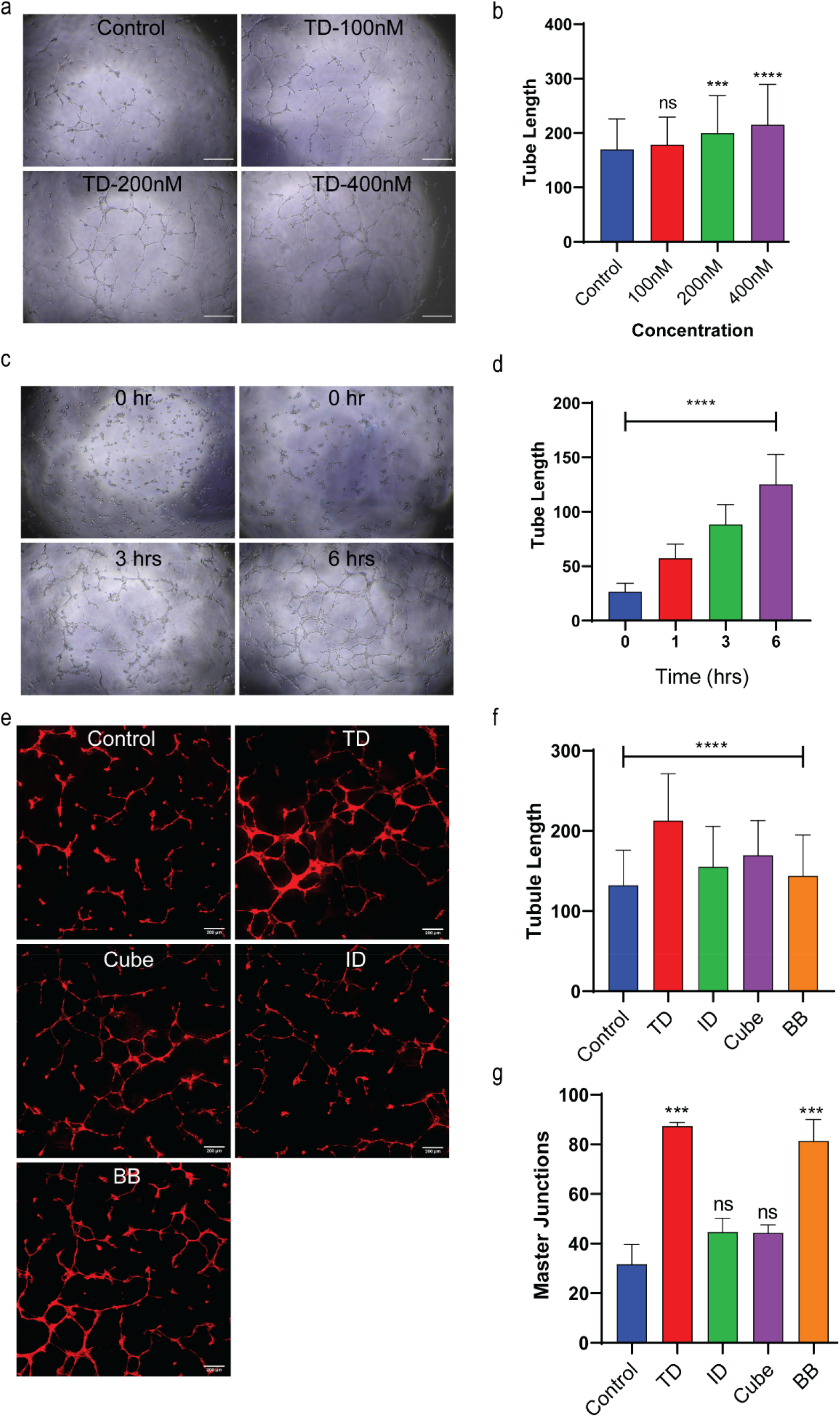
Concentration, time, and geometry-specific effect on the angiogenesis in HUVECs. **(a**,**b)** Bright field images of tubule formation at different concentration of TD (100 nM – 400 nM). Tubule formation was concentration-dependent, and longer tubules were observed at higher concentration. Tubule length was calculated from 3-4 images per condition. Error bars indicate the mean with s.d. (two-tailed unpaired t-test. **** p < 0.0001, *** p < 0.0005, ns: non-significant). **(c**,**d)** Time-dependent study to check the angiogenesis in HUVECs. Error bars indicate Standard error (ordinary one-way ANOVA, p-value < 0.0001****). **(e**,**f)** Effect of geometry on angiogenesis. Confocal images of the tubules of HUVECs stained with plasma membrane stained with Cell mask dye. DNs promoted angiogenesis with longer tubules in DNs-treated cells. Error bars indicate Standard error (ordinary one-way ANOVA, p-value < 0.0001****). **(g)** Master junctions were measured by calculating the number of junctions with 3-4 nodes and branching. Error bars indicate the mean with s.d. (two-tailed unpaired t-test. *** p < 0.0005, ns: non-significant).

Previous research has shown that tetrahedral DNA nanostructures promote the differentiation of HUVECs and this potential increases when conjugated with angiogenic peptides like VEGF^22,23^. We further checked how the geometry of DNs influences the angiogenic property of HUVECs. We tested the effect of different geometry of DNs (TD, ID, Cube and BB) on the angiogenic potential using the same protocol as before. We used 400 nM of DNs and incubated them with cells for 6 h. The tubules were stained with Cellmask^™^ in 1:1000dilution for 15mins at 37^°^C. **(Figure 3e)**. We observed that DNs had increased the tubulation of HUVECs compared to the control **(figure 3f)**. However, there was no significant difference in the tubule length among all the cages, whereas the tubules formed in control were tiny and no network was observed in control. Interestingly, the number of major junctions were more in TD and BB than ID and cube **(figure 3g)**. This trend was similar to the geometry specific bias observed in cellular uptake with HUVECs.

### DNA nanostructures promote cell migration

Cell migration refers to the movement of cells from one location to other on 2D / 3D surfaces. Cell migration is a critical process that plays a crucial role in the development, immune response, angiogenesis, cancer metastasis, and tissue regeneration^24,25^. Previous studies have reported that DNA TD enhances cell migration^26,27^. Therefore, we next studied the effect of different nanostructures on the migration of endothelial cells via scratch assay. A scratch is made on the monolayer of endothelial cells with the help of pipette tip. The influence of DNs on the migration of endothelial cells was calculated by measuring wound closure at different time points **(figure 4a)**. We observed that of all the topologies tested, BB and TD have shown more wound closure compared to the cells treated with ID and BB. The cells treated with DNs showed wound coverage by 48 h whereas; the wound was not completely covered in case of control.

**Figure 4.**
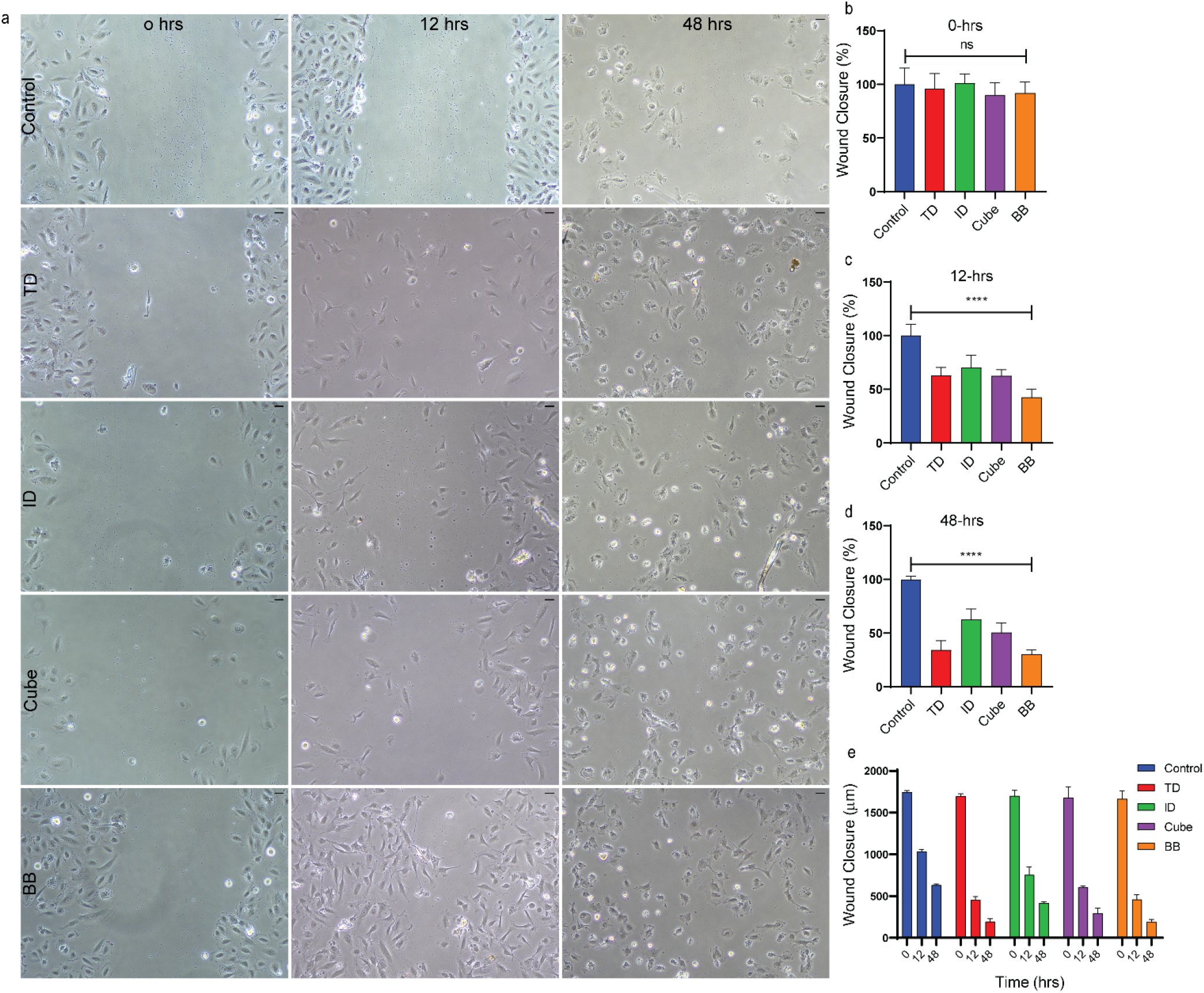
Effect of DNs on the migration of HUVECs. **(a)** Bright-field images of wound healing assay were taken to study the cell migration. The images were captured at different time points to check the wound closure. **(b)** The size of the wound was calculated at t = 0 h. **(c)** Wound closure at t = 12 h shows the migration of cells as the size of the wound has reduced. **(d)** Wound closure at t = 48 h shows the further reduction in the wound size. **(e)** Comparative quantitation of the wound closure at different time points. Error bars denote standard errors. The percent wound closure was calculated by measuring the distance of the wound. (Ordinary one-way ANOVA, P value < 0.0001****, ns: non-significant).

### Matrigel helps DNs to trigger differentiation of HUVECs

We further checked if the ECM composition had any the effect on the angiogenesis induction by DNs in HUVECs. The extracellular Matrix (ECM) plays a crucial role in endothelial cell morphogenesis. It provides structural support to cells and regulates the key signaling processes involved in angiogenesis^28^. The ECM is degraded by the proteinases and orchestrates the process of angiogenesis via the proliferation and organization of endothelial cells into multicellular 3D capillary like networks. The composition of Matrix is one of the crucial factors for tubule formation. Different products like Matrigel and geltrex mimic extracellular matrix and support the growth of the cultured cells. In order to study the effect of Matrix on the angiogenesis of HUVECs, we tested different commercially available basement membranes that mimic ECM like gelTrex; growth factor reduced Matrigel and collagen. gelTrex and Matrigel comprise of laminin, collagen-IV, entactin and heparin sulfate proteoglycan. We observed that cells were almost dead in collagen, whereas no tubule-like structures were formed in the case of gelTrex. Of all the three matrices, matrigel was the one that showed the formation of a polygonal capillary network (**Figure 5**). All the DNs triggered the tubule formation and it was statistically significant than the control untreated cells, indicating that matrix composition is indeed indispensable to observe the cellular proliferation and differentiation triggered by DNs.

**Figure 5.**
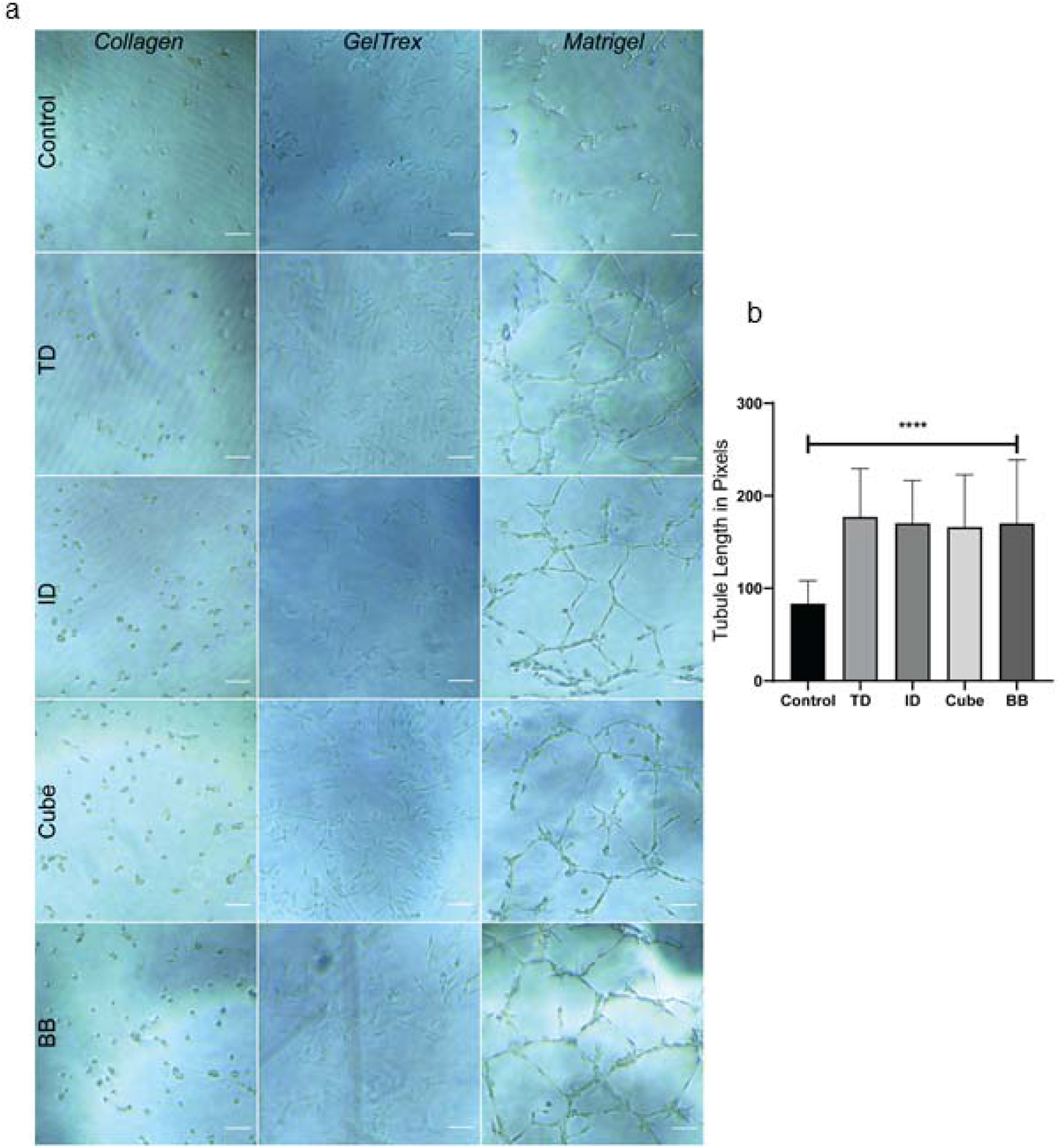
Matrix-specific differentiation of endothelial cells. (**a**) Bright field images of cells plated on different matrices to study their effect on angiogenesis or tubule formation. In case of collagen, the cells were not evenly spread. Matrigel promoted tubule formation whereas cells were undifferentiated in case of Geltrex. (**b**) Quantification of tubule length on matrigel upon treatment with different DNs. Data presented as mean±SD. Tubule length was calculated from 3-4 images (ordinary one-way ANOVA, p value < 0.0001****). Scale Bar: 100µm.

## Conclusions

Programmed differentiation of stem cells into organoids, tissues and organs is highly desired for the applications in regenerative therapies. These are the next frontiers for the cure of multiple classes of diseases like autoimmune diseases, organ failures and accidents requiring replacements. One of the key challenges in these areas are the devices or stimuli that can enhance the kinetics of differentiation of stem cells to the cell types or tissues of choice. Herein we present one such stimuli based on DNA nanodevices that could trigger cellular proliferation and differentiation. We provide a comparative assessment of different topologies of DNs and their influence on uptake, migration and differentiation of endothelial cells. Our work showed that geometry specific bias was observed during the internalization of DNs, which showed successful uptake without the aid of transfection agents. However, no angiogenesis bias was observed in the angiogenesis where all the DNs promoted differentiation compared to control. One of the key advantages of using DNA nanocages as trigger for stem cells differentiation is the well-defined 3D surface of the cages which can be easily coupled to any molecule of choice like growth factors, antigens, hormones, etc. to modulate the fate of the stem cells in a programmed way, opening up new avenues to explore for stem cells bioengineering and regenerative therapeutics.

## Materials and Methods

### 1. Materials

Human Umbilical Vein Endothelial Cells (HUVECs), Endothelial Cell Growth Basal Medium-2 (EBM) and EGM-2 SingleQuots™ Supplements were obtained from Lonza. Fetal bovine serum (FBS), PenStrap, Trypsin-EDTA (0.25%), collagen was acquired from Gibco. Phosphate buffer saline (PBS) was purchased from HyClone. The sequences used for DNA nanostructure synthesis (Table 1-4), 6X loading dye, 50 bp DNA ladder, mowiol, transferrin-A488, Hoechst and triton-X 100 were ordered from Sigma Aldrich. Nuclease free water, ammonium persulfate, ethidium bromide, TEMED, paraformaldehyde (37%) and adherent cell culture dishes were procured from Himedia. Tris-Acetate EDTA (TAE), Acrylamide/bisacrylamide sol 30% were purchased from GeNei. Magnesium chloride was ordered from SRL, India. Geltrex™ and CellMask™ were obtained from ThermoFischer. Matrigel and collagen were purchased from Corning.

**Table 1:**
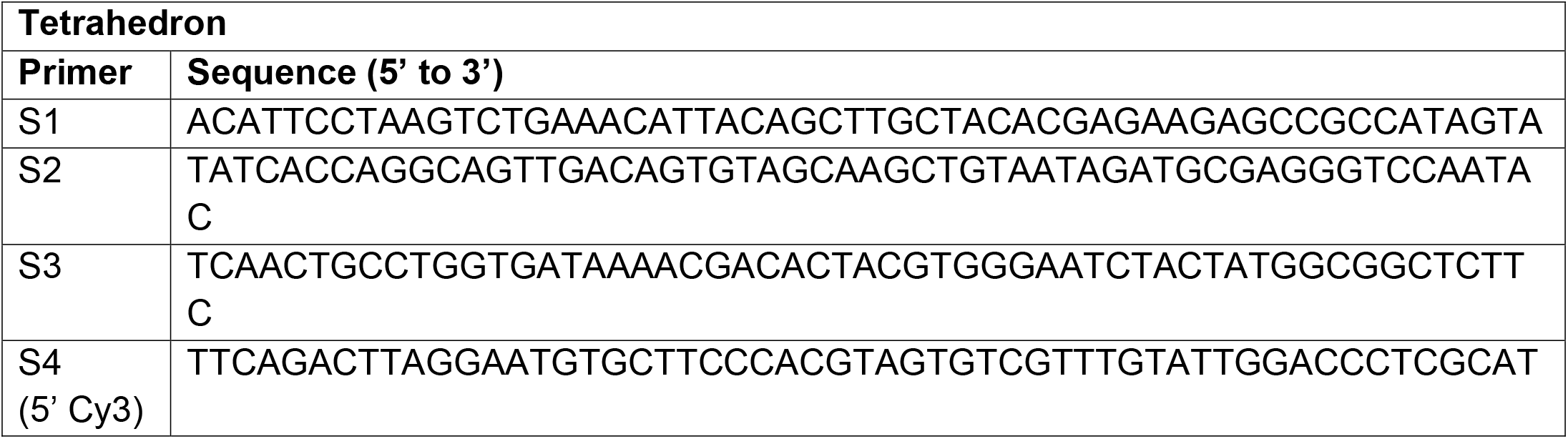
Tetrahedron.

**Table 2:**
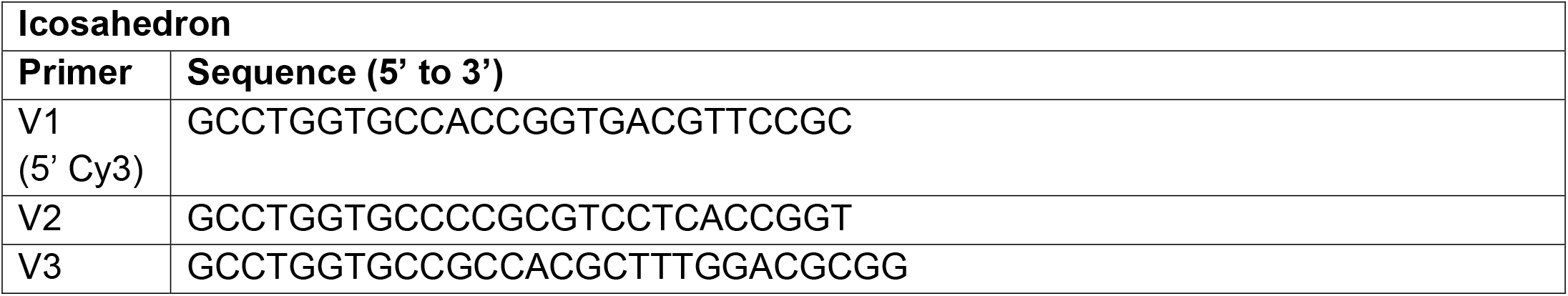
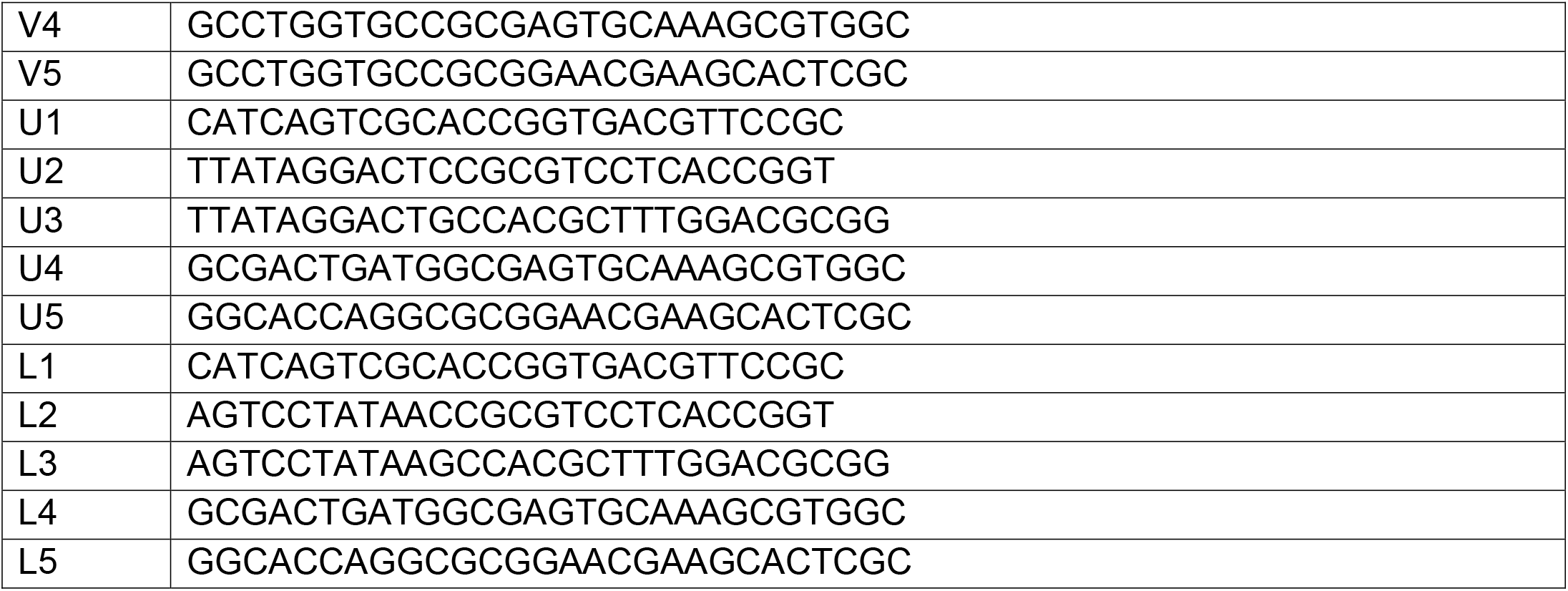
Icosahedron.

**Table 3:**
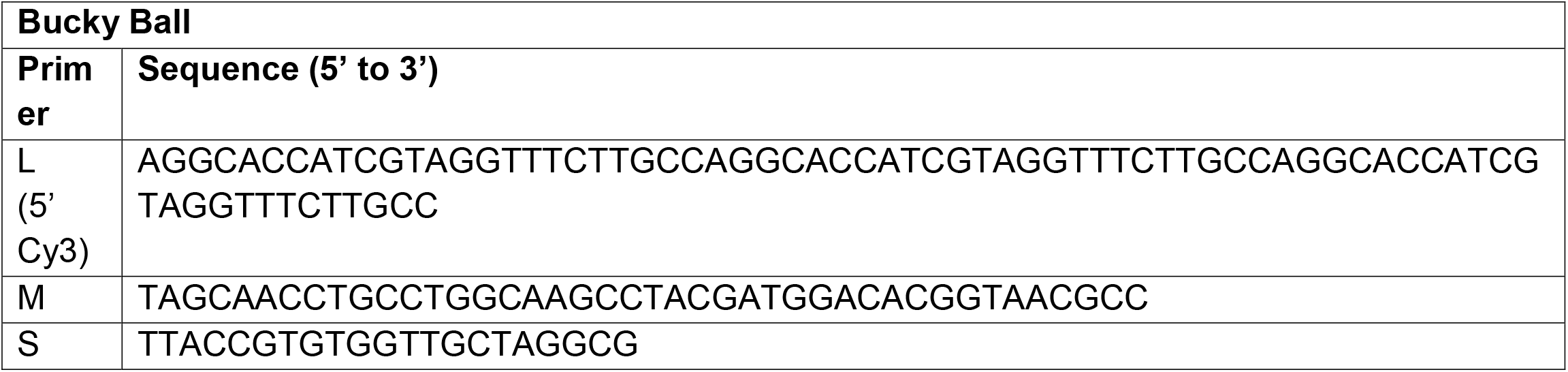
Bucky Ball.

**Table 4:**
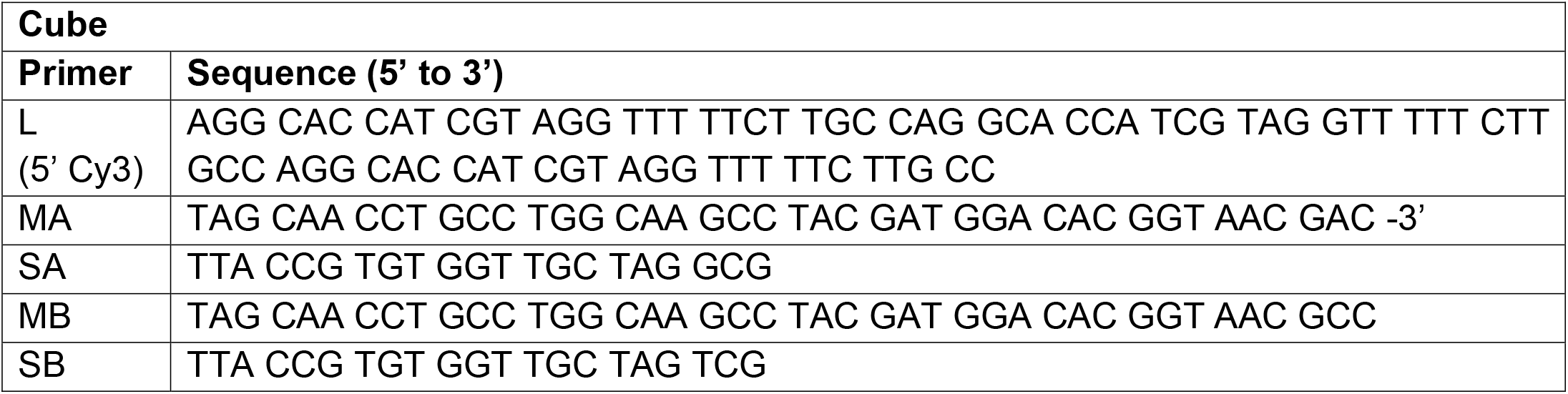
Cube.

### 2. Synthesis of DNA nanostructures

The synthesis of tetrahedron and bucky ball was done by one pot synthesis. The primers were reconstituted in nuclease free water to 100 μM stock and diluted to 10 μM working. Equimolar concentrations of four strands (S1:S2:S3:S4 - 1:1:1:1) were mixed for constructing tetrahedron; for bucky ball, the L:M:S ratio was 1:3:3. 2mM MgCl_2_ was used as catalyst. The reaction mix was heated to 95°C and annealed by decreasing the temperature by 5°C upto 4°C for 15 minutes at each step. The final concentration was 2.5 μM and 1 μM for tetrahedron and bucky ball respectively.

The icosahedron and cube were synthesized through modular assembly. For icosahedron, three 5WJ are formed in the first step (V_5_, U_5_ and L_5_) using equimolar ratio of the primers (Table 2) and 2 mM MgCl_2_. The reaction condition is similar to one pot assembly (95 to 4°C annealing). In the second step, one part of V_5_ is combined with 5 parts of U_5_ and L_5_ aided by 2 mM MgCl_2,_ resulting in two half icosahedrons (VU5 and VL5). VU5 and VL5 are assembled in 1:1 ratio in final step. The second and third step use the temperature 45°C decreasing to 4°C with 5°C decrease in each step for 4 hours at 45°C and 30 minutes for subsequent steps. For cube, the first step is creating Tile A (L:SA:MA – 1:3:3) and Tile B (L:SB:MB – 1:3:3) with 2 mM MgCl_2_. The annealing conditions are similar to one pot reaction. The second step involves 1:1 combination of Tile A and Tile B, with the reaction condition similar to second and third step of icosahedron assembly. The final concentration of icosahedron and cube were 1.6 and 1.1 μM respectively. All the DNA nanostructures were stored at 4°C until further use.

### 3. Characterization of DNA Nanostructures

Native-PAGE was used to perform Electrophoretic mobility shift assay (EMSA). 5% polyacrylamide gel was used to study the higher order structure formation. The sample preparation consisted 5 μL of DNA nanostructure with 3 μL of loading buffer and 1.5 μL of 6X loading dye. The gel was ran for 80 minutes at 80 volts. The gel was stained with EtBr stain and visualized with Gel Documentation system (Biorad ChemiDocTM MP Imaging System).

### 4. In Vitro Studies

#### 4.1. Cell Culture

HUVECs were maintained in EBM-2 supplemented with hEGF, hydrocortisone, GA-1000 (Gentamicin, Amphotericin-B), 2% FBS (fetal bovine serum), VEGF, hFGF-B, R3-IGF-1, Ascorbic Acid and Heparin. The cells were maintained at 37°C with 5% CO_2_ in a humidified incubator. The cells used in the experiments were between the passage number 3 to 7.

#### 4.2. Cellular Uptake Assay

The cells were grown up to 80-90% confluency on coverslips in 12 well plate. The cells were washed with PBS and given serum starvation for 15 minutes at 37°C. Then they were treated with 150 nM DNA nanostructure (Tetrahedron, Icosahedron, Bucky ball, Cube) and transferrin-A488 (5 μg/mL) in serum free media. They were further incubated at 37°C for 20 minutes. The cells were washed with PBS thrice and fixed with 4% PFA at 37°C for 15 minutes. They were washed with PBS to remove excess PFA and the coverslips were mounted on slides with Mowiol and Hoechst.

#### 4.3. Tubule Formation Assay

The tubule formation assay was carried out on Matrigel as basement membrane. The Matrigel was thawed at 4°C and 50 μL was used to coat each well in a 96 well plate. The plate was then left at room temperature for 10 minutes followed by 30 minutes incubation at 37°C to allow gelation. 0.4 ×10^6^ cells/mL were seeded along with 400 nM of DNA nanostructures (TD, ID, BB, Cube), and control was left without any DNA nanostructure. The cells were further imaged at different time points (0, 1, 3, 6 hours) using Nikon microscope. CellMask™ deep red was used to stain plasma membrane. Fresh working solution was prepared in 1:1000 dilution. It was added to serum free media and incubated at 37°C for 10 minutes. The images were further taken in Leica Confocal Microscope with 633 nm laser excitation. The quantification of tubules was performed in Fiji ImageJ software by calculating tubule length and counting master junctions.

#### 4.4. Cell Differentiation Assay based on Matrix

The assay procedure was similar to tubule formation assay, but three different matrixes were used to access the tubule formation. Matrigel, Geltrex™ and Collagen were used as the basement membrane.

#### 4.5. Scratch Assay

This assay was used to study cellular migration of the cells. 6-well plate was seeded and allowed to grow at 80-90% confluency. Scrape was created at the center of the wells using sterilized 200 μL tip. The wells were then gently washed with 1X PBS to remove the detached cells. They were incubated with fresh serum free media with 150 nM of DNA nanostructures (TD, ID, BB, and Cube). They were imaged at different time points (0, 6, 12, 24, 48 hours) at 10X in Nikon microscope. The images were further processed using Fiji Image J software. The distance between the scratch was measured to access the wound closure.

#### 4.6. Confocal Microscopy

The fixed slides were imaged using Confocal Scanning Laser Microscope (Leica TCS SP8). The 2D cell culture assay slides were imaged using 63x oil immersion objective. For Hoechst, 408 nm laser was used, for transferrin-A488, 488 nm laser was used. Cy3 labelled DNA nanostructures were excited using 561 nm laser. The image analysis was done using Fiji ImageJ software.

### 5. Statistical Analysis

GraphPad Prism 8.0 was used to perform all the statistical analysis. One-way analysis of variance (ANOVA) was applied and p<0.05 was considered significant. The data is represented as mean ± standard error of mean (SEM).

## Acknowledgements

We sincerely thank all the members of DB group for critically reading the manuscript and their valuable feedback. AR thank IITGN-MHRD, GoI PhD fellowship, PV acknowledges PhD fellowship from UGC-CSIR, India and IITGN for additional fellowship. DB thanks SERB, GoI for Ramanujan Fellowship, IITGN, for the startup grant, and DBT-EMR, Gujcost-DST, GSBTM, BRNS-BARC and HEFA-GoI for research grants. Imaging facilities of CIF at IIT Gandhinagar are acknowledged. Authors declare no conflict of interest.

## Table of content graphic

DNA nanocages stimulate stem cell differentiation and migration

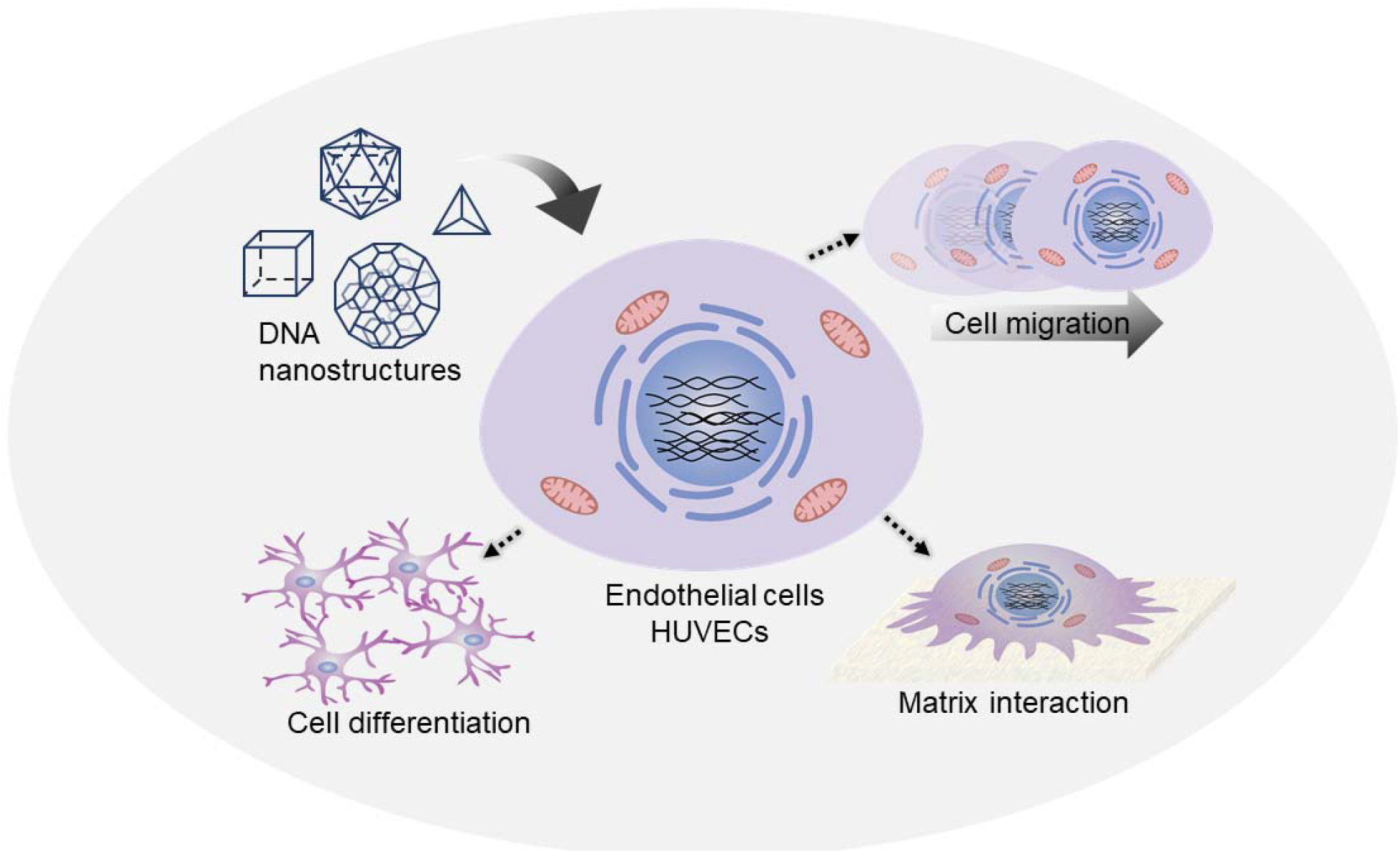

